# Transcriptional landscape of DTP-DTEP transition reveals DUSP6 as a driver of HER2 inhibitor tolerance via Neuregulin/HER3 axis

**DOI:** 10.1101/2023.06.14.544493

**Authors:** Majid Momeny, Mari Tienhaara, Deepankar Chakroborty, Mukund Sharma, Roosa Varjus, Joni Merisaari, Artur Padzik, Andreas Vogt, Ilkka Paatero, Kari J. Kurppa, Klaus Elenius, Teemu D. Laajala, Jukka Westermarck

## Abstract

The mechanisms promoting re-growth of dormant cancer cells under continuous tyrosine kinase inhibitor (TKI) therapy are poorly understood. Here we present transcriptional profiling of HER2+ breast cancer cells treated continuously with HER2 TKI (HER2i) therapy for 9 months. The data reveals specific gene regulatory programs associated with transition from dormant drug tolerant persister cells (DTPs) to proliferating DTEP (drug tolerant expanding persister) cells and eventually long-term resistance. Focusing on yet poorly understood phosphatases as determinants of therapy tolerance, expression of dual-specificity phosphatase DUSP6 was found inhibited in DTPs, but strongly induced upon re-growth of DTEPs. DUSP6 overexpression conferred apoptosis resistance whereas its pharmacological blockade prevented DTEP development under HER2i therapy. The DUSP6-driven HER2i tolerance was mediated by activation of neuregulin-HER3 axis, and consistent with the role of HER3 in widespread therapy tolerance, DUSP6 targeting also synergized with clinically used HER2i combination therapies. *In vivo,* pharmacological DUSP6 targeting induced synthetic lethal effect with HER2i in independent tumor models, and its genetic targeting reduced tumor growth in orthotopic brain metastasis model. Collectively this work provides first transcriptional landscape of DTP-DTEP transition under TKI therapy, and identify DUSP6 as a novel candidate therapy target to overcome widespread HER3-driven therapy resistance.

## Introduction

To develop therapeutic resistance, tumor cells undergo distinct evolutionary stages, starting with induction of dormancy and non-genetic drug tolerance, followed by epigenetic changes and finally resistance-conferring mutations (1–3). These different phases were originally demonstrated for EGFR-targeted therapies in non-small cell lung cancer (NSCLC) cells (4), but the concept has been expanded more recently to other malignancies including HER2+ breast cancer (1, 4–7). Based on these studies, there is ample of omics data from dormant cells (incl. DTPs) from different cancer types treated with variety of therapies. However, our understanding of the molecular mechanisms behind the regrowth of DTPs under continuous therapy is still rudimentary and to our knowledge there are no published studies describing transcriptional landscapes of DTP-DTEP transition, or transition of DTEP cells to long term resistant (LR) cells upon TKI therapies.

The human epidermal growth factor receptor 2 (HER2; encoded by *ERBB2*) is overexpressed in ∼15-25% of human breast cancers and associates with a poor patient survival (8, 9). HER2 belongs to the ERBB family of receptor tyrosine kinase (RTK) with four members: HER1 (EGFR), HER2, HER3 and HER4. Upon ligand binding, the ERBB receptors homo-and heterodimerize and activate downstream signaling pathways including PI3K/AKT and RAS/MAPK/ERK, which regulate cell proliferation, survival and the metastatic dissemination (8, 9). Multiple HER2-targeted therapies, including the monoclonal antibody trastuzumab and small molecule tyrosine kinase inhibitors (TKIs) have been approved for the treatment of HER2-overexpressing (HER2+) breast cancer (10). Application of anti-HER2 agents in combination with chemotherapy has significantly improved the patients’ outcome. However, patients initially responsive to the HER2 inhibitors (HER2is) almost inevitably succumb to disease relapse (10). Moreover, HER2+ breast tumors have an inherent tendency to develop brain metastasis, a significant clinical challenge for the treatment of these patients (11). Therefore, there is a pressing need for novel and more efficacious therapeutic strategies to overcome resistance to HER2is.

HER3 is an obligate heterodimerization partner for HER2 and plays essential roles in HER2-driven tumorigenesis, and resistance to HER2is (9, 12, 13). HER3-driven activation of PI3K/AKT and SRC are the major molecular mechanisms for resistance to HER2is (9). Consistent with the effects of HER3 overexpression, the HER3 ligand neuregulin (NRG, a.k.a Heregulin; HRG) promotes trastuzumab resistance in HER2+ breast cancer cells (9, 12). Importantly, the NRG-HER3 axis also promotes resistance to a wide range of TKIs and chemotherapies (9, 13–17). Despite the importance of HER3 in cancer progression and therapy resistance, development of HER3 small molecule inhibitors has been challenging due to its impaired kinase activity (9, 18). Moreover, the clinical activity of HER3 monoclonal antibodies either as monotherapies or in combination with chemo- and targeted therapies have been marginal (9, 19, 20). To this end, there is a pressing need to identify novel strategies to inhibit HER3 activity and/or expression for the treatment of HER3-dependent human malignancies (9, 18, 21).

There is emerging evidence that phosphatases are novel and “druggable” targets in oncology (22, 23). Inhibition of oncogenic phosphatases, or re-activation of tumor suppressor phosphatases, by small molecule therapies halt tumor growth, retard malignant progression, and enhance therapeutic sensitivity in various neoplasms (22, 23). Despite this, the contribution of phosphatases to resistance to HER2is is still poorly understood. Dual-specificity phosphatases (DUSPs) belong to the superfamily of protein tyrosine phosphatases and dephoshorylate both tyrosines and serines or threonines. A subgroup of the DUSPs are mitogen-activated protein kinase (MAPK) phosphatases that selectively interact with and dephosphorylate the MAPKs (24). For instance, DUSP6 displays a high degree of substrate selectivity for the extracellular signal-regulated kinase (ERK), but not P38 or c-Jun N-terminal kinase (JNK) (24). DUSP6 has been indicated in cancer progression, and its genetic inhibition prevents tumor cell growth (25, 26). Notably, DUSP6 is a druggable phosphatase (22, 24, 27, 28). The best characterized DUSP6 inhibitor molecule BCI (*(E)*-2-Benzylidene-3-(cyclohexylamino)-2,3-dihydro-1H-inden-1-one), is a semi-allosteric inhibitor of both DUSP1 and 6 (22, 27, 28). A more recently characterized BCI analog BCI-215 shows strict cancer cell selective cytotoxicity, overcomes leukemia therapy resistance, and selectively activates the DUSP1 and 6 target MAPKs among the 43 tested kinases (29, 30). However, the role and potential of therapeutic targeting of DUSP6 in overcoming resistance to HER2is is currently unknown.

Here we present a first transcriptional landscape of TKI treated cancer cells beyond DTP state including both DTP-DTEP and DTEP-LR transitions upon 9 months of continuous HER2i treatment. In addition to revealing global gene expression programs associated with the therapy tolerance transitions, we demonstrate that *DUSP6* has critical role in regrowth of DTEP cells under HER2i therapies. Mechanistically, DUSP6 drives non-genetic HER2i tolerance via regulation of HER3 expression and by preventing neuregulin-elicited apoptosis resistance. Collectively our findings provide a strong pre-clinical rationale to further advance in DUSP6 blockade for HER3 targeting in general, and especially for the clinical management of HER2+ breast cancer patients with resistance to HER2i.

## Results

### Development of acquired HER2i resistance by long-term treatment of drug sensitive HER2+ breast cancer cells with lapatinib

To model development of HER2i resistance from primary sensitive phase to fully resistant, the HER2+ and HER2i sensitive cell line BT474, and its brain seeking variant BT474Br (31), were exposed to therapeutically relevant 1 μM of lapatinib for 9 months (Fig. 1A). Both cell lines followed a similar pattern of lapatinib tolerance development, where at the 9-day timepoint only a few drug-tolerant persister (DTP) cells could be microscopically observed, whereas the emergence of drug-tolerant expanded persister (DTEP) population took about 6 months (Fig. 1A). Following this, the plates were fully populated by the long-term resistant (LR) cells after 9 months of continuous lapatinib treatment (Fig. 1A). Importantly, in addition to lapatinib, the LR clones of both BT474 and BT474Br displayed strong cross resistance to tucatinib (a HER2i), afatinib (a HER2/EGFR inhibitor), and neratinib (a HER2/HER4/EGFR inhibitor) (Fig. S1). This indicates that the acquired resistance is not specific to lapatinib but is driven by a mechanism that is generally relevant to the ERBB family of RTKs.

**Figure 1:**
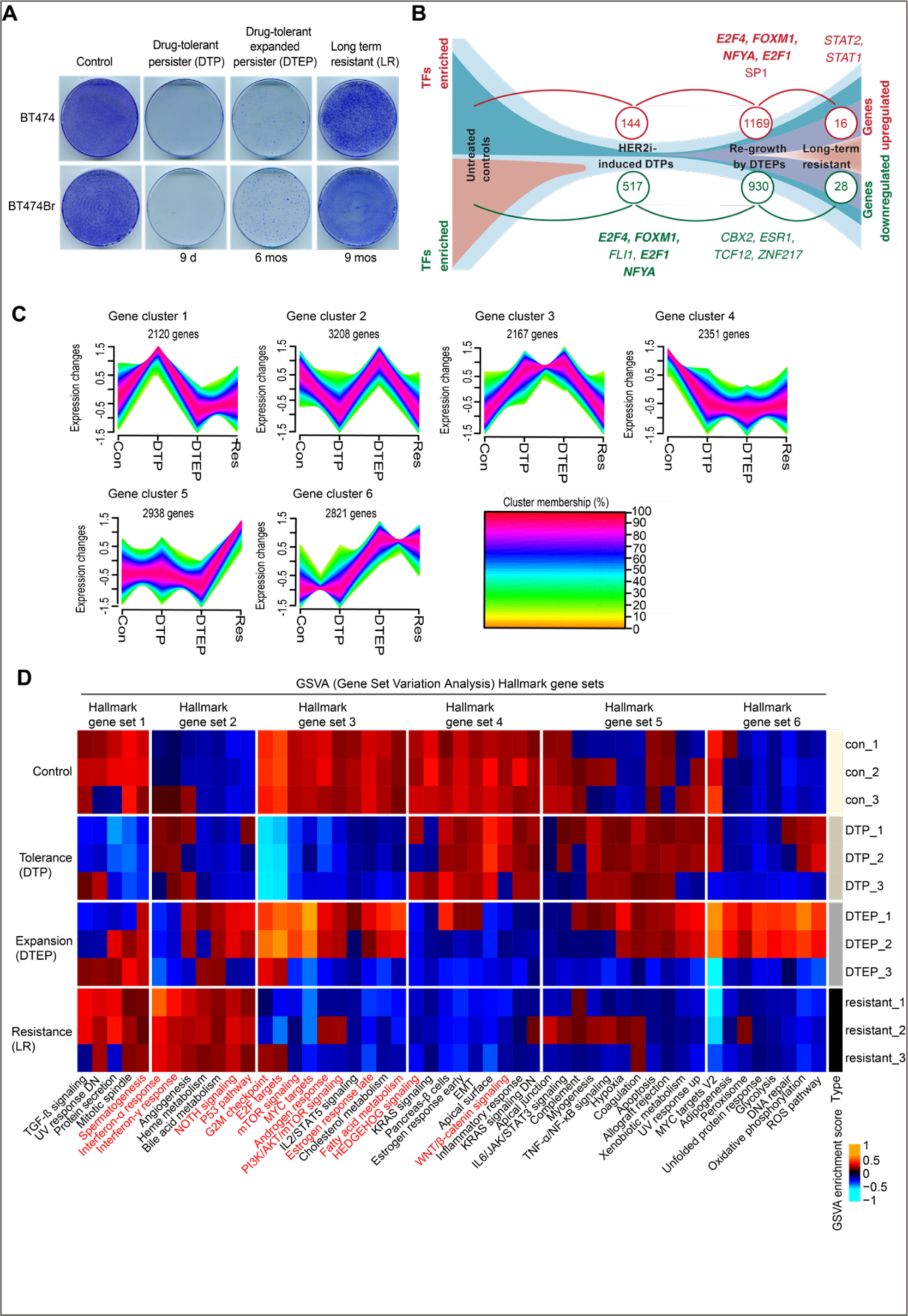
Transcriptional landscape of lapatinib tolerance and resistance development in HER2+ cells. **(A)** Development of lapatinib resistance in HER2+ breast cancer cells. BT474 and BT474Br cells were treated with 1 μM of lapatinib for 9 days (d), 6 months (mos) and 9 mos to yield DTP, DTEP and LR clones, respectively. The transcriptional profiles from each functional state of lapatinib drug tolerance and resistance development in BT474 cells were surveyed by RNA-sequencing. The cells were stained with crystal violet (0.5% w/v) and the images were acquired with an inverted microscope. **(B)** The number of the genes with a significant change in their expression during the resistance acquisition (see Table S1 for individual gene names). Transcription factor (TF) binding motifs significantly enriched in significantly regulated genes in each transition are indicated. Bolding indicates shared TF binding sites between DTP downregulated and DTEP upregulated genes. **(C)** Genes with similar patterns of expression changes during the lapatinib resistance development were clustered by unsupervised soft clustering analysis (see Table S2 for genes included in each cluster). The Cluster membership color coding indicates how tightly the genes are associated with the overall pattern of gene regulation across development of lapatinib resistance. **(D)** Differentially expressed genes and pathways were identified using the R package limma and hallmark gene sets were used for GSVA analysis to reveal signal transduction pathways involved in each step of the resistance acquisition.

### Transcriptomic landscape of acquired resistance to lapatinib

Recent studies have focused on the molecular characterization of DTP cells in response to kinase inhibitor therapies ^28,32^ but there is no published information about transcriptional profiles of TKI treated cells at the DTP-DTEP or DTEP-LR transitions. To this end, the transcriptional profiles from each functional state of lapatinib drug tolerance and resistance development in BT474 cells were surveyed by RNA-sequencing. From three technical replicates per condition, and by using statistical criteria of |logFC|>2 and Benjamini-Hochberg adjusted *p* <0.05, upregulation of 144, 1169 and 16 genes was found upon the control-DTP, DTP-DTEP, and DTEP-LR transitions, whereas the number of downregulated genes were 517, 930 and 28, respectively (Fig. 1B, Table S1). The highest number of differentially regulated genes upon the DTP-DTEP transition indicates that this is the transition phase where the cell fate is most robustly impacted during the resistance development. When assessing the patterns of gene expression changes by unsupervised soft clustering analysis ^33^, we identified six approximately similar size gene clusters with distinct regulation patterns (Fig. 1C). The genes included in these clusters are listed in the Table S2. Such regulation patterns indicate that neither gene activation nor gene repression characterize lapatinib resistance development, but unique gene expression programs are involved in each of these steps (Fig. 1B, C).

To understand gene regulatory mechanisms controlling these transitions, we predicted the transcription factor binding sites enriched on promoter regions of the differentially regulated genes (Fig. 1B, Table S3). Using FDR < 0.05 as a cut-off, the most highly enriched transcription factor elements in genes downregulated upon the control-DTP transition were E2F4, FOXM1, FLI1, E2F1 and NFYA (Fig. 1B, Table S3). Strikingly, most of these transcription factor binding sites were enriched also in the genes significantly upregulated in DTEP cells (Fig. 1B in bold, Table S3). This indicates that these transcription factors are inactivated co-ordinately upon the development of the DTP state and activated upon re-growth of the DTEPs. On the other hand, binding sites for transcription factors CBX2, ESR1, TCF12 and ZNF217 were found significantly enriched in the genes downregulated upon DTP to DTEP transition, whereas STAT1 and STAT2 were enriched among the genes upregulated in LR cells as compared to DTEPs (Fig. 1B, Table S3).

To identify cancer hallmark processes involved in each transition phase, the entire transcriptomics data was re-analysed by Gene Set Variation Analysis (GSVA), and the enriched hallmark gene sets were visualized using heatmaps with hierarchical clustering in respect to the different therapy resistance development phases. Consistent with recent evidence from other DTP models ^28-30^, MYC signaling was inhibited in the lapatinib-treated BT474 DTPs, but reactivated in DTEPs (Fig. 1D, gene sets 3 and 6). Similar gene regulation pattern was observed for E2F1 targets, G2/M checkpoint, mTOR signaling, and androgen response (Fig. 1D, gene set 3), ROS signaling, oxidative phosphorylation, and DNA repair (Fig. 1D, gene set 6). Interestingly, the DTP-DTEP transition was associated also with downregulation of several cancer hallmark gene sets, most apparently seen in set 4 where KRAS signaling, EMT, WNT/ß-catenin, and inflammatory response genes were all suppressed upon proliferation reactivation (Fig. 1D).

Collectively these results reveal the first gene expression programs associated with the DTP-DTEP transition in the TKI treated cells. The data demonstrate that unique gene clusters and biological processes are involved in each step of lapatinib resistance development. The data thus provides a rich resource for future studies of gene regulatory mechanisms in HER2i tolerance and resistance development.

### Phosphatase gene expression landscape in DTPs and DTEPs

Cancer cell phosphoproteomes are regulated by phosphatases ^15,16^, but their functional contributions to development of non-genetic kinase inhibitor therapy tolerance have been thus far poorly characterized. Therefore, we focused on the dynamics of phosphatase gene regulation during lapatinib tolerance development. Importantly, only four phosphatase genes (*CDC25A, CDC25C, DUSP6* and *SYNJ1*) were found synchronously downregulated in the DTP cells as compared to the untreated controls (Fig. 2A, C, Table S1), but upregulated in DTEPs versus DTPs (Fig. 2B, C). We rationalized that these four phosphatases might be particularly relevant for the regrowth of the DTEP cells under continuous lapatinib treatment. Even though a large group of other genes were found differentially regulated between the DTEP and LR populations (Fig. 1B, Table S1,2), none of the phosphatase genes were significantly regulated in the DTEP-LR transition (Table S1). This indicates that regulation of phosphatase gene expression could be primarily relevant to the early non-genetic phases of acquired lapatinib resistance.

**Figure 2:**
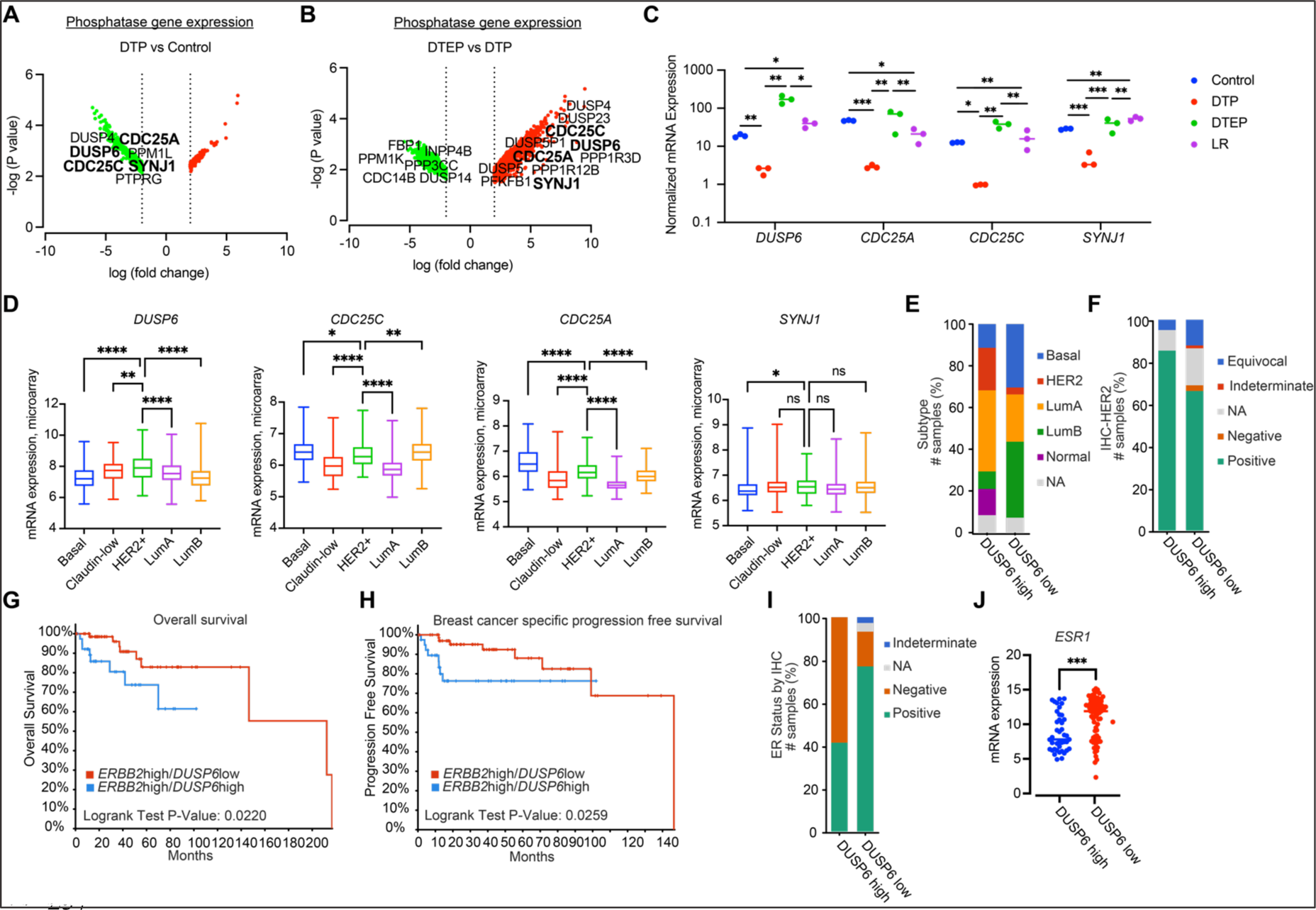
Clinical association of *DUSP6* overexpression with poor prognosis HER2+ breast cancers. **(A, B)** Volcano plot visualizing differentially expressed genes in (A) Control-DTP and (B) DTP-DTEP transitions. The volcano blots indicate all genes that were significantly regulated during these transitions (|logFC |<2 and FDR <0.05) whereas the phosphatase genes among these are indicated by names. The phosphatase genes significantly regulated in both transitions are indicated in bold. **(C)** Changes in the *DUSP6, CDC25A*, *CDC25C*, and *SYNJ1* mRNA levels during the acquisition of lapatinib resistance in BT474 cells. Data is based on RNA sequencing analysis and was analyzed by one-way ANOVA followed by Tukey’s multiple comparisons test. Statistically significant values of \**p* < 0.05, \*\*\**p* < 0.001 and ****p < 0.0001 were determined. **(D)** Differential expression of *DUSP6, CDC25A*, *CDC25C,* and *SYNJ1* in different breast cancer subtypes. Data were extracted from the METABRIC dataset and categorized into five molecular subtypes according to the PAM50 gene expression subtype classification (basal, claudin-low, HER2+, Luminal A, and Luminal B). Data were analyzed by one-way ANOVA followed by Tukey’s multiple comparisons test. Statistically significant values of \**p* < 0.05, \*\**p* < 0.01 and ****p < 0.0001 were determined. **(E)** Breast cancer patients from the TCGA-BRCA dataset were divided into DUSP6 high (LogFC>1, FDR<0.05) and low expression (LogFC<-1, FDR<0.05) groups and the clinical attributes including the subtypes were compared between the two groups. NA; not available. **(F)** Breast cancer patients from the TCGA Firehose legacy dataset were divided into DUSP6 high (LogFC>1, FDR<0.05) and low expression (LogFC<-1, FDR<0.05) groups and IHC staining positivity for HER2 was compared between the two groups. **(G, H)** Subgroup of 113 patient cases with high tumor *ERBB2* mRNA expression (LogFC>1, FDR<0.05) were divided into *DUSP6*high and *DUSP6*low groups and their overall (G) (Log-rank Test *p* value=0.0220) and disease-specific (H) survival (Log-rank Test *p* value=0.0259) was tested according to *DUSP6* status. **(I, J)** *ERBB2* high-expressing (LogFC>1, FDR<0.05) breast cancer patients from the TCGA Firehose legacy dataset were subdivided into *DUSP6*high and low groups and IHC positivity for ER (I) and the mRNA levels of *ESR1* (J) were compared between the two groups. Data were analyzed by two-tailed *t* test; \*\*\**p* < 0.001.

### Clinical association of *DUSP6* with poor prognosis HER2+ breast cancer

Among the four candidate phosphatases (*CDC25A, CDC25C, DUSP6* and *SYNJ1*) significantly regulated during both therapy tolerance transitions (Fig. 2A-C), *DUSP6* was the only one that was selectively overexpressed in HER2+ breast cancers over the other subtypes in the METABRIC dataset ^34^(Fig. 2D). The closest functional orthologue for *DUSP6, DUSP1* did not show HER2+ selective mRNA overexpression (Fig. S2A). *DUSP6* overexpression in HER2+ breast cancer was confirmed in the TCGA breast invasive carcinoma dataset ^35^(Fig. S2B). Further, when the 1082 breast invasive carcinoma samples from the TCGA BRCA-dataset ^35^ were divided to DUSP6high and DUSP6low groups based on their *DUSP6* mRNA expression levels, HER2+ and luminal A subtypes were clearly enriched among the DUSP6high samples (Fig. 2E). HER2 positivity was also found enriched among DUSP6high tumors based on HER2 immunohistochemistry of samples available for staining in the Breast Invasive Carcinoma (TCGA, Firehose legacy) dataset (Fig. 2F).

In the TCGA BRCA-dataset, patients with high *ERBB2* expression were further divided into *DUSP6*high and low groups. Comparison of the survival outcomes between the two groups indicate that high *DUSP6* expression predicts poor overall and disease-free survival among HER2+ patients (Fig. 2G, H). However, *DUSP6 m*RNA expression was not a prognostic factor in the entire patient population including the other breast cancer subtypes (Fig. S2C), further highlighting the intimate connection between DUSP6 and HER2 in breast cancer progression.

Interestingly, luminal B cancers were clearly enriched in *DUSP6*low samples as compared to *DUSP6*high (Fig. 2E). Luminal B are estrogen receptor (ER) positive cancers that often also express HER2. Clinically luminal B cancers are less aggressive than HER2+ cancers. This data may suggest that DUSP6 has a role in suppressing ER positivity, and thereby increasing the relative numbers of HER2+ cancers over luminal B cancers. Although the details of this regulation remain unclear, this hypothesis is supported by strongly decreased expression of ER protein (Fig. 2I) and *ESR1* mRNA (Fig. 2J), coding for ER, in HER2+ *DUSP6high* samples as compared to HER2+ *DUSP6low* samples. These data demonstrate that DUSP6 is clinically associated with the aggressive HER2+ breast cancer but may also have a broader role in defining breast cancer subtype development.

### DUSP6 promotes the HER2i tolerance and DTP-DTEP transition under continuous lapatinib treatment

Data above demonstrate that *DUSP6* is a clinically relevant gene in HER2+ breast cancers. As DUSP6 is also a druggable phosphatase ^15,17,20,21^, we decided to focus on its regulation and relevance in the HER2i tolerance development. To this end, we confirmed differential DUSP6 expression between different steps of the resistance acquisition in BT474 cells. Consistent with the RNA sequencing results (Fig. 2C), DUSP6 protein expression was strongly induced upon the DTP-DTEP transition (Fig. 3A). While the *DUSP6* mRNA levels were diminished in the fully resistant LR clones compared to DTEPs (Fig. 2C), its protein levels remained robustly elevated presumably via post-translational stabilization mechanisms (Fig. 3A). DUSP6 protein was also increased in BT474BrLR cells as compared to the parental cells (Fig. 3B). Notably, among the transcription factors differentially implicated upon HER2i tolerance development (Fig. 1B), forkhead box transcription factor M1 (FOXM1) binds to DUSP6 promoter (<u>ENCODE</u>). Further, coinciding with *DUSP6* expression, *FOXM1* gene expression is downregulated in DTPs and upregulated in DTEPs (Fig. 3C and Table S1). Together with its role as breast cancer oncogene involved in therapy resistance ^36^, these data imply FOXM1 as a viable candidate inducing *DUSP6* expression and proliferation during the DTP-DTEP transition.

**Figure 3:**
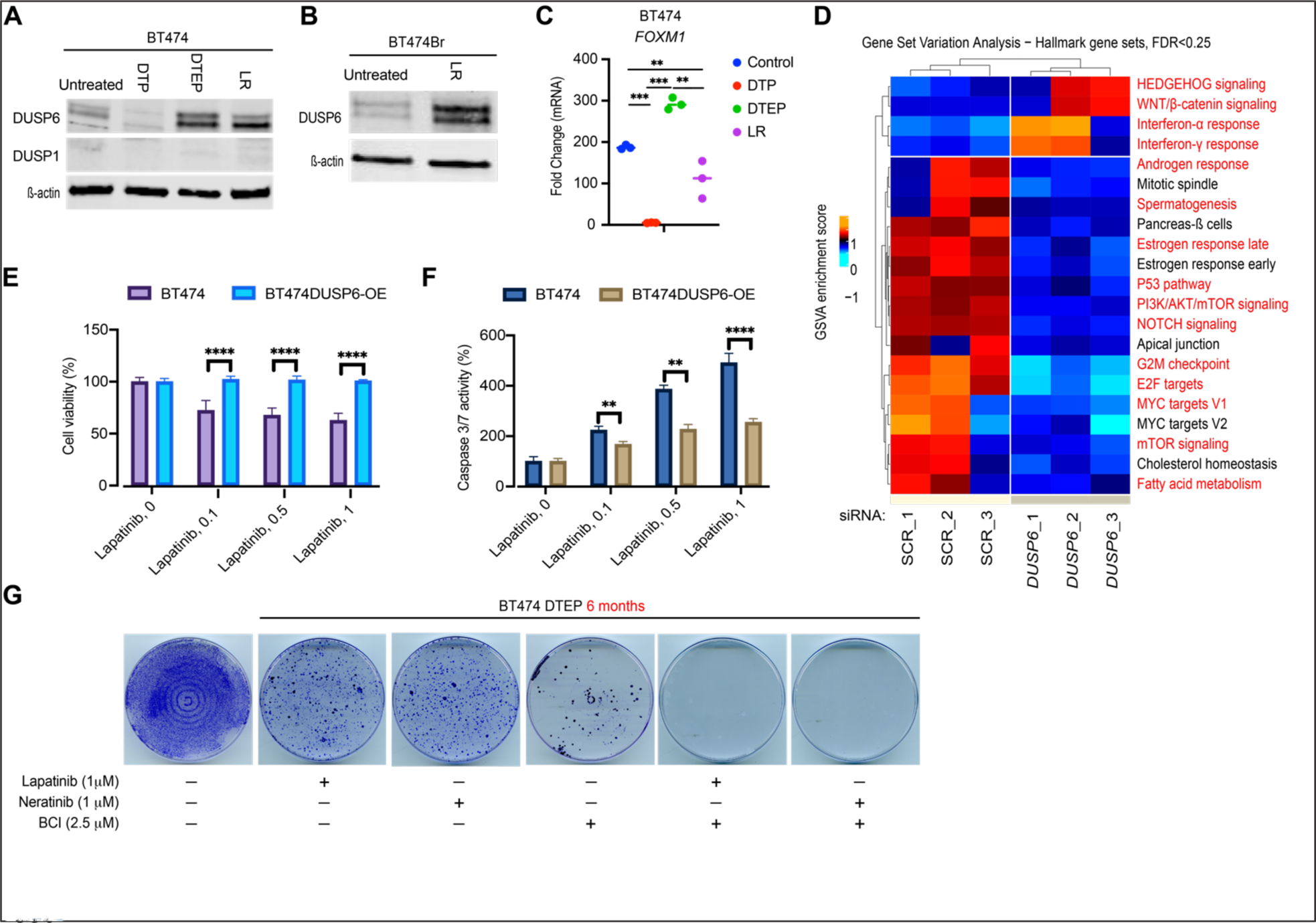
Functional involvement of DUSP6 in HER2i tolerance development. **(A)** Expression of DUSP6 protein in different stages of lapatinib resistance development by Western blot analysis. ß-actin was used as the loading control. **(B)** Increased expression of DUSP6 at LR in BT474Br cells, as shown by Western blot analysis. **(C)** Changes in the *FOXM1* mRNA levels during the acquisition of lapatinib resistance in BT474 cells. Data is based on RNA sequencing analysis and was analyzed by one-way ANOVA followed by Tukey’s multiple comparisons test. Statistically significant values of \*\*\**p* < 0.001 and ****p < 0.0001 were determined. **(D)** The transcriptional profile of MDA-MB-453 cells after *DUSP6* knockdown by 3 different siRNA was compared with 3 different scramble controls, followed by the GSVA analysis of the Hallmark gene sets. The Hallmark gene sets overlapping with the gene sets regulated to same direction in *DUSP6* low expressing DTEP cells (Fig. 1D) are indicated with red. **(E, F)** Ectopic overexpression of DUSP6 in BT474 cells inhibits lapatinib effects on cell viability (E) and apoptosis (F), as measured by WST1 cell viability assay and caspase 3/7 activity, respectively. Data were analyzed by two-way ANOVA followed by Tukey’ post hoc test. Statistically significant values of \*\**p* < 0.01 and \*\*\*\**p* < 0.0001 were determined **(G)** DUSP6 inhibitor BCI preempts DTEP development in BT474 cells treated with either lapatinib or neratinib for 6 months. The cells were stained and fixed with crystal violet in methanol (0.5% w/v) and the images were acquired with an inverted microscope.

Notably, there was a marked overlap between hallmark gene sets in DTP cells in which endogenous *DUSP6* was inhibited by lapatinib treatment and cells in which *DUSP6* was depleted by siRNA (Fig. 1D, Fig. 3D; overlapping gene sets highlighted in red). Especially interesting finding was that the *DUSP6* knockdown cells displayed a gene expression pattern linked to dormant cancer cells such as inhibition of MYC, E2F1 targets, and the PI3K/AKT/mTOR signaling, as well as activation of the interferon response ^25,28-30^. The finding that siRNA-mediated *DUSP6* depletion (Fig. 3D) recapitulates the gene expression profile in the lapatinib-induced DTP cells (Fig. 1D), clearly indicates that DUSP6 inhibition functionally contributes to the lapatinib-induced DTP state.

To ask whether the increased DUSP6 expression during the DTP-DTEP transition (Fig. 2C and 3A) functionally contributes to the survival of the DTEPs, we ectopically overexpressed DUSP6 in BT474 cells, and subjected the cells to treatment with lapatinib, neratinib, afatinib, or tucatinib. Importantly, DUSP6 overexpression was able to dampen both cell viability inhibition, and apoptosis induction, by all the tested HER2is (Fig. 3E, F, and S3A-D). As an alternative approach, BT474 cells were treated with lapatinib or neratinib alone, or in combination with small molecule DUSP6 inhibitor BCI for 6 months. Notably, as compared to the monotherapies, combination of BCI preempted the DTEP development in both lapatinib and neratinib treated cells (Fig. 3G). Strongly indicative of selective drug interaction rather than overall toxicity by BCI, the BCI used at the given concentration for 6 months did not kill all BT474 cells but potently synergized with the HER2is (Fig. 3G). The only other known BCI target, DUSP1, was neither expressed at the protein level in BT474 cells (Fig. 3A) nor its mRNA was found differentially regulated between any of the acquired resistance phases (Fig. 2A, B, Table S1). In harmony, DUSP1 overexpression had clearly weaker activity than DUSP6 overexpression in the rescue experiments (Fig. S3D). To further validate the selectivity of BCI as a DUSP6 inhibitor, we generated *DUSP6* knockout (*DUSP6* KO) MDA-MB-453 cells by CRISPR/CAS9 (Fig. S3E). The *DUSP6* KO cells were significantly resistant to BCI-elicited inhibition of cell viability, as compared to the control cells (Fig. S3F).

Collectively, these results strongly indicate that during the HER2i tolerance development, *DUSP6* downregulation is functionally relevant for establishment of the DTP phase whereas its transcriptional induction contributes to DTP-DTEP transition.

### DUSP6 targeting kills HER2i resistant breast cancer cells and synergizes with HER2i combination therapies

After discovering the role for DUSP6 in the development of HER2i tolerance, we wanted to address its role in HER2+ breast cancer cells with stable *de novo* or acquired HER2i resistance. To this end, we compared pharmacological DUSP6 targeting against a library of available small molecule modulators of phosphatases ^15,16^, or inhibitors of oncogenic signaling pathways and anti-apoptotic proteins. Across either *de novo* (MDA-MB-453), or acquired (BT474LR and BT474BrLR) HER2i resistant cells lines, inhibition of cell viability by phosphatase targeting was observed only with DUSP6 inhibitor BCI, its derivative BCI-215, and with FTY-720 that reactivates protein phosphatase 2A (PP2A) by SET inhibition ^37^ (Fig. 4A). Based on the response to kinase inhibitors and anti-apoptotic antagonists, these HER2i resistant, but BCI sensitive cell lines were co-dependent on PI3K/AKT, PLK1 and AXL kinase activities, and on the anti-apoptotic proteins IkB, survivin, cIAP and/or XIAP (Fig. 4A). Corroborating the selective role for DUSP6 as the anti-apoptotic target of BCI and BCI-215, siRNA-mediated depletion of *DUSP6*, but not *DUSP1*, induced apoptosis in MDA-MB-453 cells (Fig. 4B). *DUSP6* depletion also induced apoptosis in another HER2i resistant HER2+ cell line MDA-MB-361 (Fig. S4A). Furthermore, the CRISPR/CAS9 targeted MDA-MB-453 *DUSP6* KO cells showed severely impaired long-term colony growth potential (Fig. 4C). These results indicate that among druggable human phosphatases, DUSP6 is a therapeutic target candidate for HER2i resistant HER2+ breast cancer.

**Figure 4:**
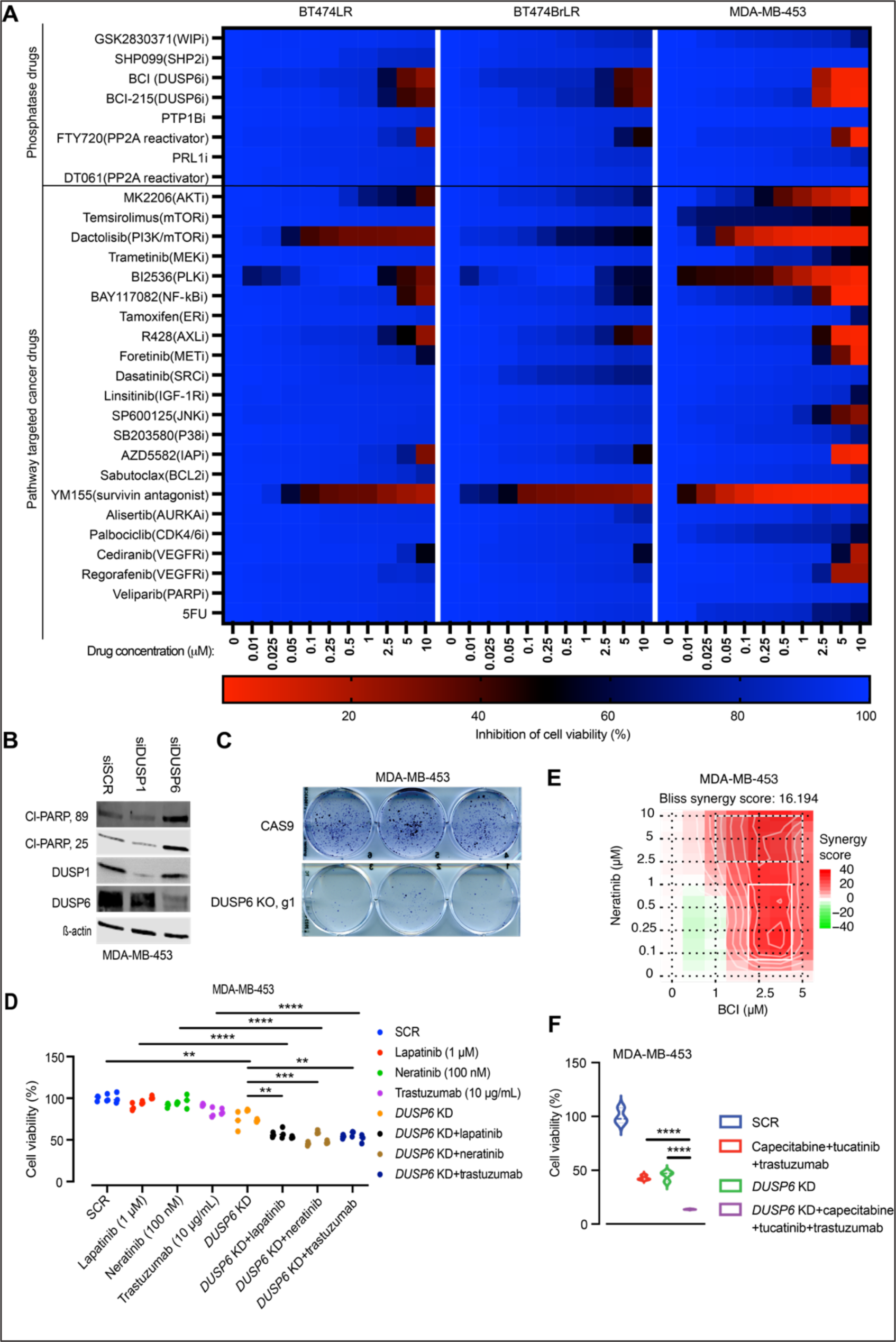
Both genetic and pharmacological DUSP6 targeting overcomes HER2i resistance. **(A)** The anti-proliferative activities of a library of small molecule modulators of phosphatases, kinases and anti-apoptotic proteins in the *de novo* and acquired HER2i resistant cells. The cells were treated with the increasing concentrations (in μM) of the compounds for 48 h and cell proliferation was measured using WST-1 assay. **(B)** RNAi-mediated *DUSP6* knockdown, but not *DUSP1*, induces apoptotic cell death in MDA-MB-453 cells, as shown by PARP-1 cleavage. The blots are representative of three independent experiments with similar outcomes. **(C)** *DUSP6* knockout hinders the clonogenic survival of MDA-MB-453 cells, as compared to the CAS9 controls. The cells were seeded at low density and maintained for 10 d. The colonies were stained/fixed with 0.5% crystal violet in methanol and imaged using an inverted microscope. **(D)** *DUSP6* siRNA knockdown increases sensitivity of HER2i resistant MDA-MB-453 cells to HER2-targeted therapies. Cell growth was measured by WST-1 cell viability assay after 48 h of drug treatment. Data were analyzed by one-way ANOVA followed by Tukey’s multiple comparisons test. Statistically significant values of \*\**p* < 0.01, \*\*\**p* < 0.001 and \*\*\*\**p* < 0.0001 were determined. **(E)** A 2D synergy map of neratinib-BCI combination in MDA-MB-453 cells calculated by Bliss SynergyFinder ^40^. The cultures were treated with increasing concentrations of the compounds for 48 h and cell viability was measured by WST-1 cell viability assay. **(F)** *DUSP6* siRNA knockdown increases sensitivity of HER2i resistant MDA-MB-453 cells to combination with capecitabine and two HER2 targeting therapies, trastuzumab and tucatinib. Cell growth was measured by WST-1 cell viability assay after 48 h of drug treatment. Data were analyzed by one-way ANOVA followed by Tukey’s multiple comparisons test. Statistically significant values of \*\**p* < 0.01, \*\*\**p* < 0.001 and \*\*\*\**p* < 0.0001 were determined.

Notably, the genetic *DUSP6* targeting sensitized the HER2i resistant MDA-MB-453 cells to several HER2 targeting approaches. Indeed, whereas the parental MDA-MB-453 cells were resistant to the clinically relevant concentrations of lapatinib, neratinib and trastuzumab, these HER2is regained their therapeutic activity in the *DUSP6-* siRNA targeted cells (Fig. 4D, S4B). The impact of genetic *DUSP6* targeting in HER2i sensitization was recapitulated by BCI treatment across primarily resistant cell lines (Fig. 4E and S4C). Specifically, whereas neratinib did not impact HER2i resistant cell survival at 100 nM concentration (Fig. 4E), BCI greatly synergized with these low micromolar doses of neratinib (Fig. 4E; white rectangular, and S4C). Additionally, *DUSP6* depletion increased sensitivity to the combination of neratinib and capecitabine (Fig. S4D), which improves progression-free survival (PFS), and time to the CNS disease intervention in HER2+ breast cancer ^38^. Furthermore, *DUSP6* knockdown enhanced sensitivity to the combination of tucatinib+trastuzumab+capecitabine (Fig. 4F), which improves PFS and overall survival in patients with HER2+ metastatic breast cancer ^39^.

Together with DUSP6 overexpression experiments in HER2i sensitive cells (Fig. 3E, F), these data provide strong evidence that DUSP6 contributes to the HER2i resistance both in monotherapy and combination therapy settings. The results also indicate that pharmacological inhibition of DUSP6 may be an efficient therapeutic approach to target HER2i resistance in HER2+ cells.

### DUSP6 inhibition overcomes HER2 inhibitor resistance *in vivo*

To validate the *in vivo* relevance of the results, we used both genetic and pharmacological targeting of DUSP6 in HER2i resistant xenograft models. To start with, we evaluated the impact of CRISPR/CAS9-mediated *DUSP6* knockout on the xenograft growth of MDA-MB-453 cells in immunocompromised BALB/cOlaHsd-Foxn1nu mice. Importantly, the two *DUSP6* KO clones showed indistinguishable antitumor effects as compared to the control cells (Fig. 5A). After obtaining this genetic evidence that DUSP6 promotes HER2+ cell tumor growth, we asked if pharmacological DUSP6 blockade overcomes HER2i resistance *in vivo*. For this purpose, the mice with MDA-MB-453 or HCC1954 tumor size ∼100 mm^3^ were randomized into four treatment groups; vehicle, lapatinib/neratinib (50 mg/kg), BCI (50 mg/kg), and lapatinib/neratinib+BCI. Importantly, validating the *in vivo* HER2i resistance of both chosen HER2+ cell models, tumors from these cell lines were fully resistant to clinically relevant doses of either lapatinib or neratinib (Fig. 5B-E). Notably, both MDA-MB-453 and HCC1954 tumors also displayed strong resistance to BCI monotherapy (Fig. 5B-E), indicating tumor microenvironment-mediated impact as compared to the *in vitro* cultures. However, combination of BCI with either lapatinib or neratinib cancelled the HER2i resistance phenotype very efficiently (Fig. 5B-E). Indicating for potential clinical utility, DUSP6 and HER2-targeted therapies displayed a clear synthetic lethal drug interaction when assessed by a waterfall blot (Fig. 5C, E). The dramatic combinatorial activity of DUSP6 and HER2 targeting was also evidenced by total acellularity of the MDA-MB-453 xenograft tumor after 24 d of treatment and based on hematoxillin and eosin (H&E) staining (Fig. 5F). Consistent with previous studies with BCI ^18,19,41,42^, we did not observe any apparent signs of toxicity or weight loss (Fig. S4E) in mice treated with either monotherapies, or with neratinib plus BCI combination.

**Figure 5:**
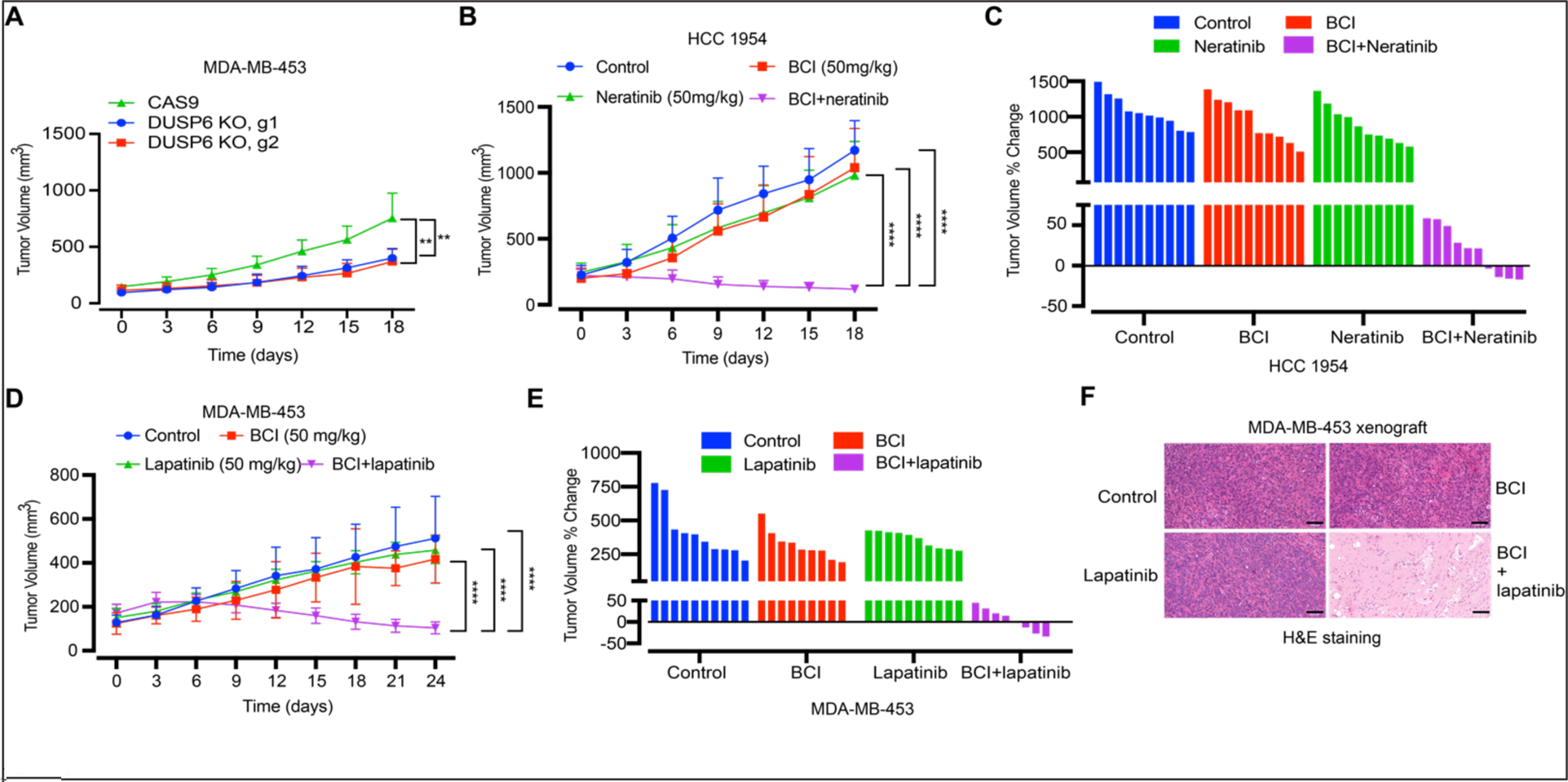
DUSP6 inhibition overcomes HER2 inhibitor resistance *in vivo*. **(A)** *DUSP6* knockout decreases tumor growth in a xenograft model of MDA-MB-453 cells. **(B-E)** The effect of BCI in combination with lapatinib and neratinib in xenograft models of HCC1954 (3 x 10^6^ cells) and MDA-MB-453 (5 x 10^6^ cells), respectively. Mice with tumor size ∼100 mm^3^ were randomized into the experimental and the control groups and tumor volumes were measured every 3 d. Data were analyzed by one-way ANOVA followed by Tukey’s multiple comparisons test. Statistically significant values of **p < 0.01 and ****p < 0.0001 were determined. **(F)** H&E staining of the MDA-MB-453 xenograft tumors from the control, lapatinib, BCI and lapatinib+BCI groups at day 24. Scale bar 200 μm.

### DUSP6 is a superior target over AKT in targeting HER2i resistant cells

The PI3K/AKT/mTOR signaling pathway is a key HER2 downstream effector ^43^ and its constitutive activity is strongly associated with HER2 therapy resistance ^44-46^. In this vein, clinical studies of HER2i plus PI3K/AKT inhibitors in patients with HER2+ breast cancer with progression on HER2i-based therapy demonstrated some clinical activity, but did not lead to approval of these combinations ^47,48^. On the other hand, PI3K/AKT activation was one of the DUSP6-driven hallmark gene sets associated with the DTP-DTEP transition (Fig. 1D and 3E) whereas the DUSP6-dependent cell models were co-dependent on the PI3K/AKT activity (Fig. 4A). Therefore, it was important to compare the quantitative and qualitative differences between DUSP6 blockade versus direct AKT inhibition in combination with the HER2i in the resistant models. Having demonstrated phenocopying growth effects between genetic *DUSP6* inhibition and BCI (Fig. 4), as well as resistance of *DUSP* KO cells to BCI (Fig. S3F), these experiments were mostly performed by comparing the compounds MK2206 (AKTi) and BCI (DUSP6i) as alternative pharmacological HER2i combination approaches.

Both BCI and MK2206 synergized with already low micromolar concentrations of lapatinib and neratinib in MDA-MB-453 and HCC1954 cells, respectively (Fig. 6A, B, S4C and S5A). However, even though DUSP6 and AKT inhibition were quantitatively comparable in their synergistic cell viability effects, DUSP6 targeting had qualitatively clearly superior pro-apoptotic activity. Regardless of efficient inhibition of AKT phosphorylation, MK2206 did not induce apoptosis alone, or in combination with lapatinib (Fig. 6C, D). In contrast, treatment with BCI plus lapatinib triggered apoptosis across all the tested cell models (Fig. 6C, D, and Fig. S5B).

**Figure 6:**
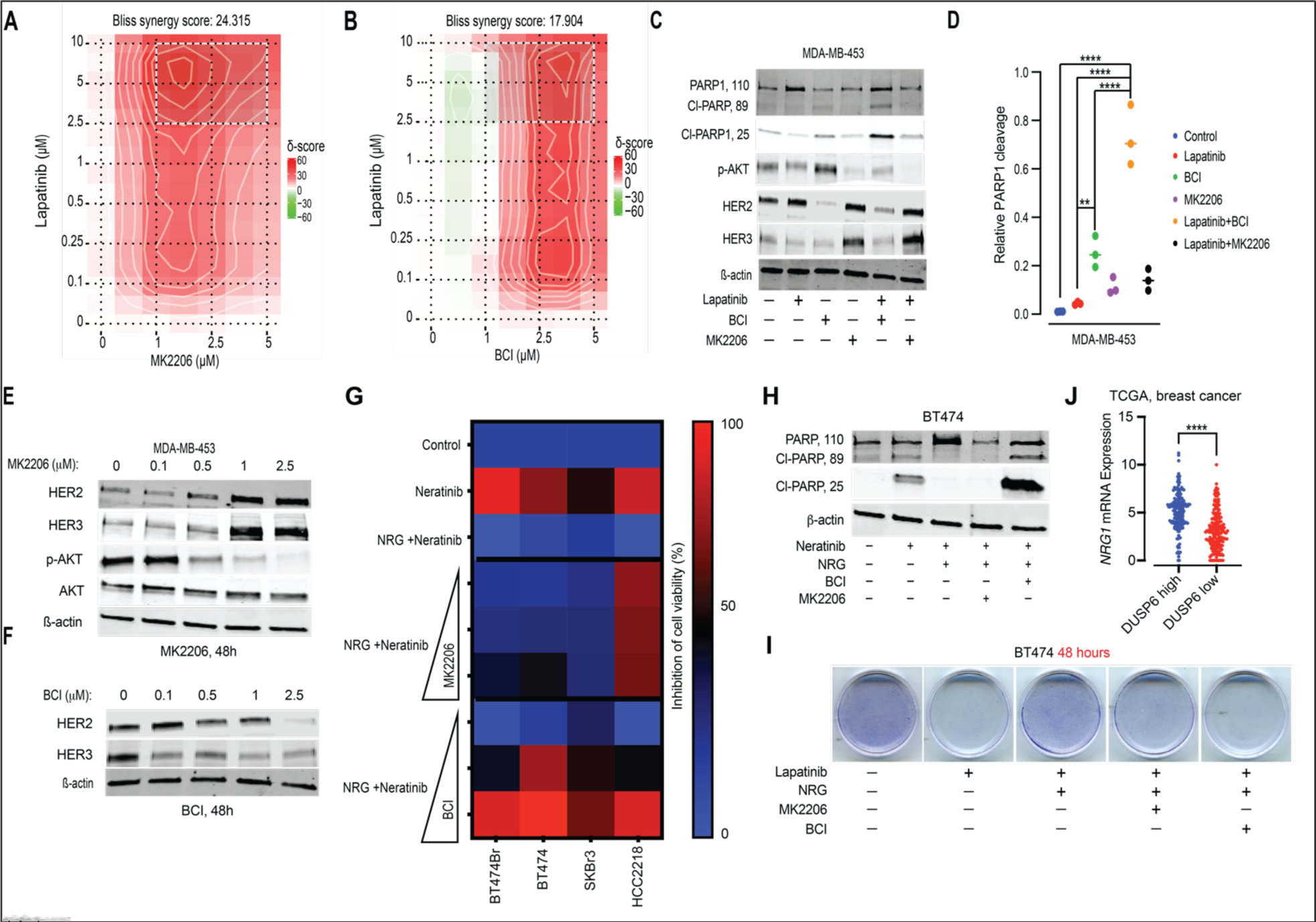
DUSP6 is a superior combination therapy target over AKT in targeting HER2i resistant cells. **(A)** A 2D synergy map of lapatinib-MK2206 or **(B)** lapatinib-BCI combination in MDA-MB-453 cells calculated by the Bliss SynergyFinder. The cultures were treated with increasing concentrations of the compounds for 48 h and cell viability was measured by WST-1 cell viability assay. **(C)** Comparison of the effects of lapatinib+BCI and lapatinib+MK2206 on HER2, HER3 and cleaved PARP-1 protein levels by Western blot analysis. The cells were treated with lapatinib (1 μM), MK2206 (2.5 µM) and BCI (2.5 µM) for 48 h. The blots are representative of three independent experiments with similar outcomes. **(D)** Quantification of PARP1 cleavage from three repeats of (C). Data were analyzed by one-way ANOVA followed by Tukey’s multiple comparisons test. Statistically significant values of **p < 0.01 and ****p < 0.0001 were determined. **(E)** The dose-dependent effects of MK2206 and **(F)** BCI on the expression of HER2 and HER3 protein levels in MDA-MB-453 cells by Western blot analysis. The blots are representative of three independent experiments with similar outcomes. **(G)** Comparison of the effects of neratinib+BCI and neratinib+MK2206 on NRG-mediated rescue from the anti-proliferative activity of neratinib. The cells were treated with NRG (10 ng/mL), lapatinib (1 µM), MK2206 (1, 2.5, and 5 µM), and BCI (1, 2.5, and 5 µM) for 48 h and cell viability was measured by WST-1 assay. **(H)** Comparison of the effects of neratinib+BCI and neratinib+MK2206 on NRG-mediated evasion from neratinib-induced apoptotic cell death, as measured by Western blot analysis for cleaved PARP-1. The cells were treated with NRG (10 ng/mL), lapatinib (1 μM), MK2206 (2.5 µM) and BCI (2.5 µM) for 48 h. The blots are representative of three independent experiments with similar results. **(I)** Comparison of the effects of lapatinib+BCI and lapatinib+MK2206 on NRG-mediated rescue from the anti-growth activity of lapatinib, as shown by crystal violet staining. The cells were treated with NRG (10 ng/mL), lapatinib (1 μM), MK2206 (2.5 µM) and BCI (2.5 µM) for 48 h, stained/fixed with 0.5% crystal violet in methanol and imaged by an inverted microscope (images acquired at 10x magnification). **(J)** Breast cancer patients from the TCGA-BRCA dataset were divided into *DUSP6* high (LogFC>1, FDR<0.05) and low expression (LogFC<-1, FDR<0.05) profiles and the *NRG1* mRNA levels were compared between the two groups. Data were analyzed by two-tailed t test; \*\*\*\**p* < 0.0001.

### DUSP6 targeting overcomes NRG-mediated HER2i therapy tolerance

HER3 is a key pro-survival receptor in HER2+ breast cancer cells ^2,5^. Further, compensatory induction of HER3 is thought to be one of the primary mechanisms behind the resistance to the combination of HER2 inhibition and AKT targeting ^6,49^. Accordingly, HER3 induction by AKT inhibition was observed in MDA-MBA-453 cells treated with either MK2206, or PI3K inhibitor GDC-0084 alone, and with MK2206-lapatinib combination (Fig. 6C, E and Fig. S6A). In contrast, BCI decreased HER2 and HER3 expression as a monotherapy, and in combination with lapatinib (Fig. 6C, F). Directly linking HER3 regulation to apoptosis, RNAi-mediated *HER3* knockdown triggered PARP1 cleavage in MDA-MB-453 cells (Fig. S6B). These findings indicate that lack of compensatory induction of HER3 may explain apoptosis induction in BCI+lapatinib treated cells, as compared to MK2206+lapatinib combination (Fig. 6C, D).

Tumor microenvironment derived HER3 ligand NRG drives resistance to HER2is via a paracrine mode ^50,51^. With this background, we further compared DUSP6 and AKT in the NRG-induced rescue of the HER2i in sensitive cell lines. As expected, ^50,51^, NRG treatment induced tolerance to both neratinib and lapatinib across all four tested primarily sensitive cell lines (Fig. 6G and Fig. S6C). The NRG-mediated rescue from both HER2is was abrogated in a concentration dependent manner after treatment with BCI, but not with MK2206 (Fig. 6G and Fig. S6C). The only exception was HCC2218 cells, in which also MK2206 reversed NRG-elicited HER2i tolerance (Fig. 6G and Fig. S6C). These results were validated at the level of differential apoptosis inducing capacity between DUSP6 and AKT targeting. Whereas NRG completely prevented HER2i-elicited induction of apoptosis in BT474 cells, and AKT inhibition could not reverse this, DUSP6 inhibition fully restored apoptotic activity (Fig. 6H). To link these results to development of HER2i tolerance development, we further demonstrated that targeting DUSP6, but not AKT, reversed the NRG-elicited effects on lapatinib tolerance in the DTP assay (Fig. 6I). We further confirmed that the effects of BCI on the reversal of NRG-elicited rescue were due to DUSP6 inhibition, as siRNA-mediated *DUSP6* depletion also abrogated the NRG-induced cell survival in BT474 cells (Fig. S6D). Finally, providing clinical validation for the link between NRG and DUSP6, *NRG* mRNA was significantly overexpressed in breast tumors with high *DUSP6* mRNA expression (Fig. 6J). Importantly, deregulation of NRG-ERBB axis was evidenced also during the DTP-DTEP transition of lapatinib treated cells as, concomitantly with *DUSP6* regulation (Fig. 2C), *NRG3* and *ERBB4* were found upregulated in DTEP versus DTP cells (Fig. S6E).

Collectively these results demonstrate that DUSP6 inhibition is superior to AKT blockade as a HER2i combination therapy strategy due to its capacity to overcome the NRG-elicited survival signaling, and trigger apoptosis.

### A DUSP6-HER3 feed forward loop drives the HER2i tolerance

The results above indicate that DUSP6 may drive HER2i resistance by promoting the expression of HER3. Accordingly, knockdown of *DUSP6* reduced HER2 and HER3 protein levels in MDA-MB-453, HCC1954, and the HER3+ triple negative breast cancer cell line MDA-MB-468 (Fig. 7A and S7A, B). Notably, clinically high *DUSP6* tumors had significantly higher expression of tyrosine 1298 phosphorylated HER3 (Fig. 7B). Indicative of transcriptional regulation, *DUSP6* inhibition decreased *HER2* and *HER3* mRNA expression, but did not impact HER2 and HER3 protein stability (Fig. S7C, D). Further, indicative of selective effects on HER2 and HER3 among the ERBB receptors, EGFR expression was unaffected by DUSP6 targeting in MDA-MB-468 (Fig. S7B). Notably, both HER2 and HER3 were downregulated 3 h after BCI treatment, and this coincided with the induction of ERK phosphorylation and destabilization of DUSP6, both serving as clear signs for BCI target engagement (Fig. 7C). The tumor material from the BCI-treated MDA-MB-453 xenograft model was further used for *in vivo* validation of HER2 and HER3 as DUSP6 downstream targets. Indeed, when evaluated by IHC, BCI-treated tumors displayed vastly decreased tumor cell immunopositivity for HER3 and to lesser extent HER2 (Fig. 7D), and without effects on the overall tumor cellularity (Fig. 5F).

**Fig. 7.**
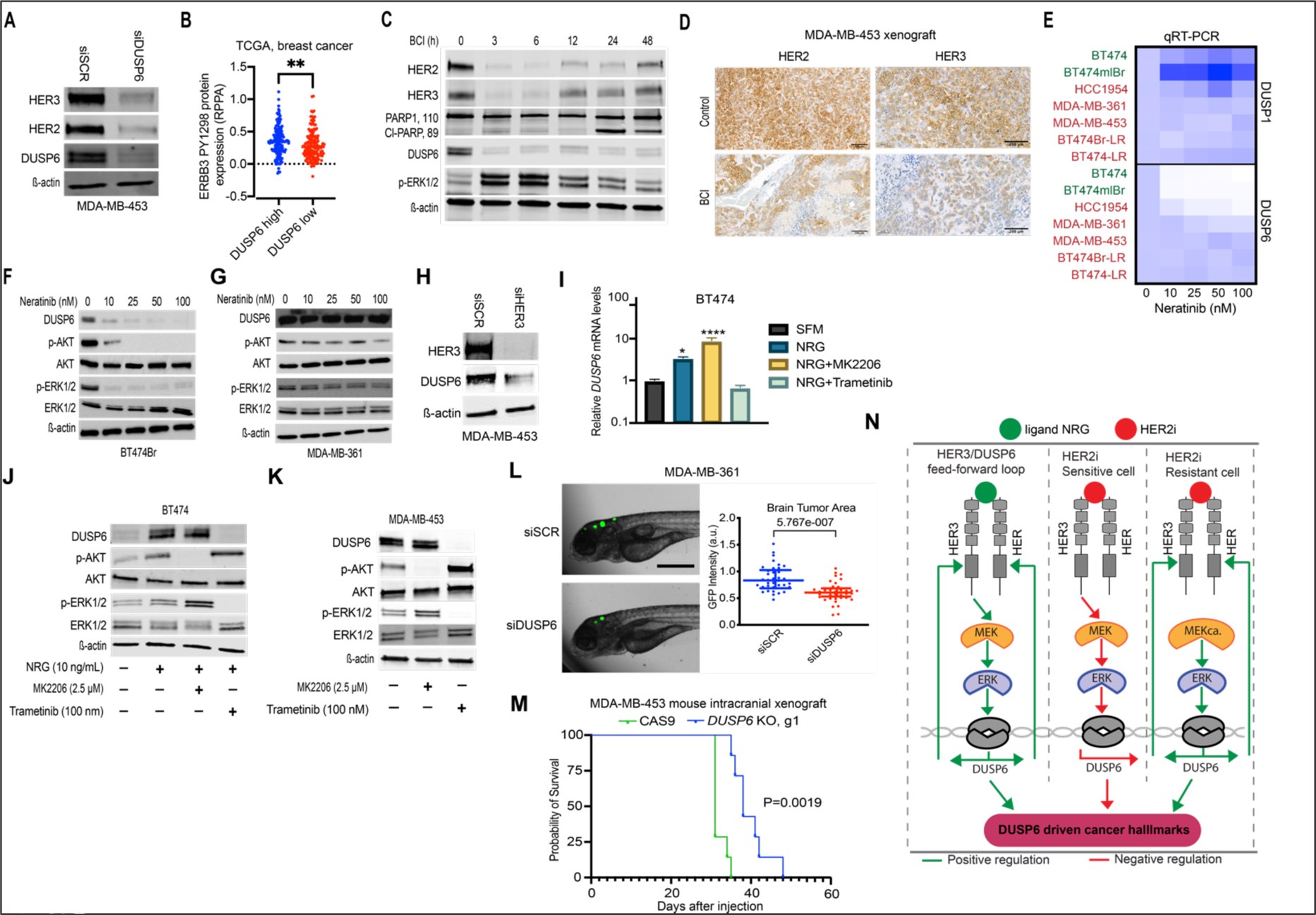
Identification of HER3-DUSP6 positive feedback loop. **(A)** *DUSP6* knockdown inhibits HER2 and HER3 protein expression in MDA-MB-453 cells as shown by Western blot analysis. The blots are representative of three independent experiments with similar results. **(B)** Breast cancer patients from the TCGA-BRCA dataset were divided into *DUSP6* high (LogFC>1, FDR<0.05) and low expression (LogFC<-1, FDR<0.05) profiles and the p-HER3^Y1298^ levels were compared between the two groups. Data were analyzed by two-tailed t test; \*\**p* < 0.01. **(C)** Effect of BCI (2.5 μM) on HER2 and HER3 expression and PARP1 cleavage in MDA-MB-453 cells. The blots are representative of three independent experiments with similar results. DUSP6 inhibition and ERK phosphorylation serve as protein markers for BCI target engagement to DUSP6. **(D)** Effects of BCI (50 mg/kg) therapy on HER2 and HER3 protein levels in the MDA-MB-453 xenograft tissue at day 24. The samples are adjacent from those shown in Fig. 5F. Scale bar 200 µm. **(E)** *DUSP1* and *DUSP6* mRNA levels were determined by qRT-PCR analysis after treatment with increasing concentrations of neratinib for 48 h in indicated cell lines. Red denotes for HER2i resistant cell lines and green HER2i sensitive cells. **(F, G)** The effect of treatment with increasing concentrations of neratinib for 48 h on DUSP6, p-AKT and p-ERK1/2 was measured by Western blot analysis in BT474Br and MDA-MB-361 cells, respectively. The blots are representative of three independent experiments with similar outcomes. **(H)** The effect of *HER3* knockdown on DUSP6 expression in MDA-MB-453 cells was determined by Western blot analysis. The blot is representative of three independent experiments with similar results. **(I)** NRG-mediated induction of *DUSP6* mRNA via MEK activation as measured by qRT-PCR analysis after treatment with NRG (10 ng/mL), MK2206 (2.5 µM), and trametinib (100 nM) for 48 h. Data were analyzed by one-way ANOVA followed by Tukey’s multiple comparisons test. Statistically significant values of \**p* < 0.05 and \*\*\*\**p* < 0.0001 were determined. (**J)** The effect of NRG on DUSP6 expression BT474 cells via MEK activation. The cells were serum-starved for 24 h, followed by treatment with NRG (10 ng/mL), MK2206 (2.5 µM) and trametinib (100 nM) for 48 h. The blots are representative of three independent experiments with similar outcomes. **(K)** Trametinib inhibits DUSP6 expression in HER2i resistant MDA-MB-453 cells through MEK blockade. The cells were treated with MK2206 (2.5 µM) or trametinib (100 nM) for 48 h. The blots are representative of three independent experiments with similar results. **(L)** The effect of *DUSP6* knockdown on the brain metastatic outgrowth of MDA-MB-361 cells in a zebrafish model. Data were analyzed by two-tailed *t* test; \*\*\*\**p* < 0.0001. Scale bar 100 µm. **(M)** *DUSP6* knockout improves overall survival in an intracranial murine model as compared to the control. Survival data were analyzed by log-rank Mantel–Cox test, \*\**p* < 0.01. **(N)** A schematic illustration of the HER3/DUSP6 feed forward loop in HER2+ breast cancer cells. NRG binding to HER3 induces MEK/ERK-mediated *DUSP6* expression which feeds back to increased HER2 and HER3 expression (left panel). In HER2i sensitive cells (middle panel) inhibition of HER3 results in *DUSP6* inhibition and loss of DUSP6 driven cancer hallmarks. In HER2i resistant cells (right panel), MEK is not inhibited by HER2i but its constitutive activity (MEKca.) drives DUSP6-HER2/3 positive feed-back loop resulting in HER3-mediated multitherapy resistance and cancer progression.

After identifying previously unrecognized regulation of HER3 by DUSP6, we asked whether the HER3 reciprocally regulates DUSP6 expression. Indeed, *DUSP6* mRNA expression was potently inhibited already with the smallest tested concentration of neratinib or lapatinib in the HER2i sensitive cells (BT474, BT474Br) (Fig. 7E, Fig. S7E, in green). However, among the HER2i resistant cell lines, *DUSP6* was not inhibited by HER2is in MDA-MB-361, MDA-MB-453, BT474Br-LR, and BT474-LR cells, whereas HCC1954 showed an intermediate phenotype (Fig. 7E, Fig. S7E, in red). Additionally, this regulation was specific to *DUSP6,* as *DUSP1* mRNA expression was not inhibited by HER2 targeting in any of the tested cell modes (Fig. 7E, Fig. S7E). Clinically *DUSP6* and *HER3* mRNA expression also correlated in HER2+ cancer samples in the TCGA-BRCA dataset ^52^(Fig. S7F). Differential regulation of DUSP6 expression by HER2i in sensitive (BT474Br, BT474) versus resistant (MDA-MB-361, MDA-MB-453, BT474BrLR) cells was validated at the protein level by western blotting (Fig. 7F, G and S8A, B). Importantly, RNAi-mediated knockdown of *HER3* decreased DUSP6 expression (Fig. 7H, I). HER3 dependent regulation was further supported by induction of DUSP6 mRNA and protein expression by treatment of serum-starved BT474 or BT474Br cells with the HER3 ligand NRG (Fig. 7I, J and S8C, D). Induction of DUSP6 was selective for NRG, since hepatocyte growth factor (HGF), which activates MET RTK, failed to increase DUSP6 in BT474 cells (Fig. S7I, J). To gain insights into the mechanism by which HER3 regulates DUSP6, the NRG treated cells were co-treated either with MK2206, or the MEK1/2 inhibitor trametinib. Consistent with a previous report that DUSP6 is a transcriptional ERK1/2 target ^17^, treatment with trametinib inhibited both ERK phosphorylation and DUSP6 expression in all tested cell lines (Fig. 7I-K and Fig. S8E), whereas AKT inhibition by MK2206 did not impact DUSP6 expression (Fig. 7I-K). Notably, while MEK blockade inhibited DUSP6 expression in the HER2i resistant cells (Fig. 7K and S8E), resistance to HER2 inhibition in DUSP6 regulation correlated with the capability of HER2i to impact ERK phosphorylation (Fig. 7F, G and S8A, B). This indicates that the failure of HER2i to inhibit DUSP6 in acquired resistant cells is due to resistance mechanism upstream of MEK.

Based on the newly discovered role for DUSP6 in promoting HER3 expression, combined with the key role for HER3 in brain metastasis from HER2+ breast cancer ^53,54^, we decided to investigate contribution of DUSP6 in the brain metastatic outgrowth. To this end, we quantitated the zebrafish intracranial tumor area derived from GFP+ MDA-MB-361 cells transfected prior tumor implantation either with control or *DUSP6* targeting siRNA. As shown in Fig. 7L, the zebrafish larva with *DUSP6* targeted cells had significantly smaller tumors 3d after the intracranial injection. Further, mice bearing the intracranial *DUSP6* KO MDA-MB-453 tumors had a significantly longer survival time as compared with the CAS9 control group (Fig. 7M).

Together with the NRG rescue experiments (Fig. 6G-I), these results strongly indicate that inhibition of the newly discovered DUSP6-HER3 feed-forward loop has a major contribution in HER2i response (Fig. 7N; left and middle panels). On the other hand, lack of HER2i-elicited *DUSP6* inhibition in the resistant models (Fig. 7E, G and Fig. S8B) provides a novel explanation for HER2i resistance in these cells (Fig. 7N; right panel). Considering the important role of the NRG-HER3 axis in mediating cancer therapy resistance across different types of therapies ^2,6-10^, this newly identified role of DUSP6 in promoting HER3 expression may explain the cross-therapy resistance observed in TKI treated cells ^6,25,27^. Collectively, these data validate DUSP6 as a novel therapy target candidate to overcome HER2i resistance, and to inhibit growth of brain metastasis in HER2+ breast cancer.

## Discussion

Cancer therapy resistance is initiated by non-genetic signaling rewiring resulting in major changes in the epigenetic and transcriptional landscapes of DTPs ^25,27,28^. It is commonly believed that transition from dormant DTPs to proliferating DTEPs is a pivotal step towards development of the disseminated disease, and eventually late-stage genetic changes which determine the ultimate therapy resistance ^25,26^. Here, we report the first transcriptomic analysis of the DTP-DTEP transition in TKI treated cancer cells. In addition to importance of the discovery of the role of DUSP6 in DTP-DTEP transition, the transcriptome data will provide a rich resource for future studies to understand different stages of the HER2i resistance development. Based on the fuzzy clustering data (Fig. 1C and Table S2), the gene clusters 1-4 that display significant differences between the parental and the DTP cells could include additional targets relevant for the establishment of dormant stage of lapatinib-treated DTPs. On the other hand, the gene clusters 1, 2 and 6, contain genes that are strongly regulated upon the DTP-DTEP transition and are therefore likely to be involved in the evasion of the DTP cells from lapatinib-induced growth inhibition. These longitudinal transcriptomics profiles also allowed us to identify potential master transcription factors involved in the lapatinib resistance development. These hypotheses, and the role of the differentially expressed genes associated with each transition during lapatinib resistance development need to be validated in future studies stemming from this novel resource.

Phosphatases have recently emerged as novel druggable targets for cancer therapy ^15-17^. However, their contribution to development of non-genetic TKI therapy tolerance is still poorly understood. In the present study, we asked which phosphatases might contribute to the development of HER2i tolerance and resistance, and explored their potential role as therapy targets in combination with the HER2i therapies in HER2+ breast cancer. Via transcriptomics analysis of lapatinib tolerance development over six months, we discovered an important role for DUSP6 in the re-growth of HER2i tolerant cells and showed that its genetic and pharmacological blockade potentiates HER2i sensitivity *in vitro* and *in vivo*. Our findings further illustrate DUSP6-mediated regulation of HER2 and 3 as a novel mechanism for its oncogenic activity. While demonstrated here for the first time in the context of HER2i therapy resistance, genetic DUSP6 inhibition has recently been shown to inhibit malignant phenotypes in other cancer types ^17-19^. Consistent with these reports, genetic inhibition of *DUSP6* in our study resulted in significant inhibition of HER2+ breast cancer cell viability, HER2i resistance, colony forming potential, and *in vivo* tumor growth. Mechanistically, DUSP6 overexpression blocked DUSP6 inhibition converted cytostatic response to HER2i to an apoptotic response which is considered as a paramount for cancer therapy strategies aiming for cancer cure. On the other hand, overexpression of DUSP6 converted HER2i sensitive cells to resistant in both cell viability and apoptosis assays. Importantly, all our main conclusions remained valid regardless of whether DUSP6 was inhibited either by siRNA, CRISPR/CAS9, or by pharmacological inhibitors BCI and BCI-215. We further demonstrated nearly immediate target engagement (Fig. 7C), as well as relative resistance of *DUSP6* knock-out cells to BCI-elicited cell viability inhibition (Fig. S2F). Most importantly, the results identified DUSP6 as a novel combination therapy target with existing clinical HER2 targeting strategies including chemotherapy combinations (Fig. 4D-F, 6B-D). In that regard, our xenograft results demonstrate significant potential for pharmacological DUSP6 inhibition in overcoming HER2i resistance *in vivo* by using two different HER2i resistant cell lines (MDA-MB-453 and HCC1954), and two different HER2is (lapatinib and neratinib). Whereas neither HER2 nor DUSP6 inhibition alone did not have therapeutic effect on tumors, the combination showed very potent synthetic lethal phenotype further validating our *in vitro* results. The lack of monotherapy effect of BCI, combined with clear evidence for downstream target mechanism engagement *in vivo* (Fig. 7D), also alleviates concerns about overall cellular toxicity behind the BCI-elicited HER2i synergy. This is consistent with previous findings demonstrating antitumor effects with BCI and BCI-215 without obvious systemic toxicity in *in vivo* models of gastric cancer, leukemia, and malignant peripheral nerve sheath tumor ^18,19,41,42^.

Dysregulation of the PI3K/AKT/mTOR pathway due to *PIK3CA* mutations or *PTEN* loss drives HER2i therapy resistance ^44-46^. Despite this, combination of the PI3K/mTOR or AKT inhibitors with HER2i-containing regimens has shown marginal therapeutic efficacy in clinical trials ^55-57^. These disappointing results are explained at least partly by the release of negative feedback from mTOR and AKT on HER2 and 3 and their activation by NRG ^49,58^. Here we identify DUSP6 targeting as a novel approach to target NRG-HER3 axis and thereby overcome HER2i resistance. Furthermore, our data demonstrated that DUSP6 inhibition, but not AKT inhibition, impairs NRG-induced rescue from the anti-proliferative activity of HER2is. These are important advantages favoring DUSP6 blockade versus AKT inhibition as a future strategy for treatment of HER2+ breast cancer.

A particularly important translational finding in our study is that DUSP6 defines therapy sensitivity of the lapatinib resistant cells in response to several ERBB TKIs with different specificities, and to their chemotherapy combinations (Fig. 4D-F, S4D). This is consistent with the proposed HER3-DUSP6 positive feedback mechanism, as HER3 or NRG overexpression confer resistance to several cancer drugs ^2,6-10^. Thereby identification of DUSP6 targeting as a novel approach for HER3 inhibition may have broad ramifications in different combination therapy settings across different cancer types. HER2 and 3 play also essential roles in the development of brain metastasis which causes death of more than 30% of stage IV HER2+ breast cancer patients ^4,54,59^. To this end, DUSP6-mediated regulation of HER2 and 3 suggested that its inhibition may retard development of brain metastasis. Indeed, our findings directly support these conclusions as *DUSP6* knockout in tumor cells improves survival of mice with intracranial HER2+ tumors and the outgrowth of HER2+ breast tumor cells in a zebrafish intracranial model. Linking our results further to future therapy of HER2+ brain metastasis, we showed that *DUSP6* knockdown increases sensitivity to tucatinib+trastuzumab+capecitabine combination regimen (Fig. 4F), which show significant clinical activity in HER2+ breast cancer patients with brain metastasis ^39^.

In summary, we provide first transcriptional map of DTP-DTEP transition under TKI tolerance development in cancer. The results specifically identify DUSP6 targeting as a novel approach to target HER3-mediated ERBB TKI resistance. Ultimately, the work provides proof-of-principle evidence to encourage development of next generation DUSP6 inhibitors (with brain penetrance) to test the clinical relevance of the presented therapy scenarios.

## Materials and methods

### Reagents

Tissue culture reagents including regular RPMI, DMEM, RPMI and FBS were purchased from Sigma. The recombinant NRG, HGF and β-estradiol (E2) were from Peprotech. BCI was purchased from Axon Medchem. BCI-215 was provided by Dr. Andreas Vogt, University of Pittsburgh Drug Discovery Institute, Pittsburgh, PA, USA. The HER2 inhibitors and the other compounds used in the drug screening were purchased from Adooq Bioscience. All the agents were dissolved in DMSO and the final concentration of DMSO did not exceed 0.1% [v/v] in all the treatments.

### Cloning and plasmids

pBABE-puro-gateway-ERBB2 was a gift from Matthew Meyerson (Addgene plasmid No. 40978; http://n2t.net/addgene:40978; RRID:Addgene_40978). ERBB3 wild-type was cloned from pBABE-puro-gateway-ERBB3 ^60^ into pLenti CMV Puro DEST (w118-1), a gift from Eric Campeau and Paul Kaufman (Addgene plasmid # 17452;http://n2t.net/addgene:17452; RRID:Addgene_17452) through Gateway cloning ^61^ to create pLenti-CMV-Puro-ERBB3.

### Generation of stable lines

To overexpress wild-type ERBB2, pBABE-puro-gateway-ERBB2 was transfected (using Fugene6 transfection reagent; Promega Catalog # E2692) into amphotropic Phoenix HEK293T cells (a gift from Dr. Garry Nolan) to generate retroviruses, which were used to transduce MDA-MB-453, as described previously ^61^. pLenti-CMV-Puro-ERBB3 was co-transfected with virus-packaging plasmids pMLDg/pRRE (addgene #12251), pMD2.G (addgene #12259), and pRSV-Rev (addgene #12253) into HEK293T cells using Fugene6 transfection reagent to produce lentiviruses. The lentivirus-containing supernatant was used to transduce MDA-MB-453 to over-express wild-type ERBB3. After viral transduction cells were treated with 1 µg/mL puromycin (Gibco) for 48 h to select the cells with stable expression of the respective introduced transgenes.

For DUSP1 and 6 overexpression, lentiviral particles containing full length of either DUSP1 (Genecopoeia) or DUSP6 (Addgene #27975) or control empty (Genecopoeia) vector were generated in HEK293FT packaging cell line (complete medium: high glucose DMEM, 10% FBS, 0,1mM NEAA, 1mM MEM Sodium Pyruvate, 6mM L-Glutamine, 1% Pen/Strep and 0,5mg/ml Geneticin) by transient transfection of transfer vector 2^nd^ generation packaging plasmid-psPAX2 (Addgene #12259) and envelope vector-pMD2 (Addgene #12260) with the ratio (7:2:1) using calcium-phosphate precipitation method. Seventy-two hours post-transfection medium containing viral vectors was collected, concentrated for 2 h by ultracentrifugation in swing-out rotor SW-32Ti (Beckman Coulter), 26,000g, resuspended in residual medium and flash-frozen in liquid nitrogen. Functional titer ̴1×10^8^ was measured in HEK293FT cells and FACS (BD LSRFortessa, Becton Dickinson). To obtain stable overexpression of DUSP1, DUSP6 or double DUSP1+6 population on day zero, 8×10^4^ cells were seeded in a 24-well plate. 24 h later, the cells were transduced with MOI 60 of lentiviral stocks in a low volume of full media. Medium containing viral particles was removed 16 h later. Cells expressing DUSP1 and GFP indicative of lentiviral integration were collected by fluorescence assisted cell sorting (BD FACSaria II cell sorter, Becton Dickinson). FACS gating was set at 10% top high fluorescence signal. DUSP6 transduced BT474 cells were selected with 3 µg/mL of puromycin. DUSP1 and 6 expressing cells were obtained by sequential transduction (MOI 2, 6, 10), puromycin selection and later GFP fluorescence assisted cell sorting (BD FACSaria II cell sorter, Becton Dickinson). The levels of protein were confirmed by Western blot analysis.

### FACS

MDA-MB-453 cells transduced with lentiviruses encoding wild-type ERBB3 were washed with azide-free PBS, trypsinized and suspended in ice-cold sorting buffer (PBS +1% Goat serum, Life techonolgies catalog # PCN5000). The cells were incubated with anti-ERBB3 (MAB3481, R&D systems) for 1 h on ice, and with anti-mouse Alexa Fluor 405 (A-31553, Invitrogen) for 30 min on ice in a dark environment. Single cell suspension was analyzed and sorted on Sony SH800 Cell Sorter to select cell-pools with high surface ERBB3 expression.

### CRISPR/CAS9 knockout system

A two-component CRISPR system was used to generate *DUSP* KO cells ^62^. DUSP6 sgRNAs (seq#1-CATCGAGTCGGCCATCAACG, seq#2-GACTGGAACGAGAATACGGG, seq#3-CCATGATAGATACGCTCAGA) were selected using DeskGEN platform and cloned according to F. Zhang lab protocol. Separate lentivectors containing spCas9 (lentiCas9-Blast a gift from Feng Zhang (Addgene plasmid # 52962) and sgRNA (lentiGuide-Puro a gift from Feng Zhang (Addgene plasmid # 52963) were produced in HEK293FT packaging cell line by transient cotransfection. Shortly, 40-70% confluent HEK293FT cells were used for transfections with 14 μg of transfer vector, 4 μg of packaging vector psPAX2 (gift from Didier Trono (Addgene plasmid # 12260), 2 μg envelope vector pMD2.G (gift from Didier Trono (Addgene plasmid # 12259) mixed in 0.45 mL water, 2.5M CaCl2, and 2x HeBS (274 mM NaCl, 10 mM KCl, 1.4 mM Na2HPO4, 15 mM D-glucose, 42 mM Hepes, pH 7.06) per 10 cm dish. Before adding to the cells, the DNA-HeBS mix was incubated for 30 min at room temperature. After overnight incubation medium with DNA precipitate was gently removed from the cells and replaced with a full fresh medium. Media containing viral particles was collected after 72 h, spun at 300 rpm for 5 min at room temperature to remove cell debris, filtered through 0.45 µm PES filter, and concentrated by ultracentrifugation for 2 hours at 25,000rpm, 4°C (Beckman Coulter). The pellet containing lentiviral particles was suspended in the residual medium, incubated for ∼2 hours in +4°C with occasional mild vortex, aliquoted, snap-frozen and stored in −70°C. P24 ELISA measured physical lentiviral titer with a serial dilution of virus stock according to manufacturer protocol.

To generate DUSP6−/− clones in MDA-MB-453, 1e+05 cells were seeded on a 24-well plate. The next day the cells were transduced with Lenti-Cas9 (MOI 1, 5, 10), and 72 h later, 8 μg/mL of Blasticidin was applied to select only Cas9 expressing cells. Cells transduced with the smallest number of Lenti-Cas9 particles that survived after the parallel control well was cleared proceeded to the next step. In the second stage, the mixed pool of stably expressing Cas9 cells was transduced with Lenti sgRNA vectors (MOI 5,10,15,20), and 72 h later, 1 μg/uL Puromycin was applied on the cells to select double-positive Cas9+/Lenti sgRNA+ cells. Based on Western blot results, cell populations showing the highest reduction in DUSP6 protein levels were single sorted (Sony SH800 cell sorter, Sony Biotechnology Inc) and re-grown into a clonal cell population. On average, about 20 clones per sgRNA population were screened using Western blotting and qPCR. Sanger sequencing was used to confirm full knockout status.

### Cell Culture and Transfections

All cell lines were purchased from the American Type Culture Collection (ATCC) and were maintained at 37 °C and 5% CO_2_ in a humidified incubator and cultured according to the ATCC recommendations. Cell line authentication was performed by STR profiling and using Cell ID^TM^ system (Promega). The cultures were routinely tested for mycoplasma contamination. The information about siRNAs for *DUSP1*, *DUSP6*, *HER2*, *HER3*, *AKT1*, *BIRC5* and negative control siRNAs are in Table S4. Transient transfections were performed with lipofectamine RNAiMAX reagent (ThermoFisher) according to the manufacturer’s instructions.

### Cell growth assays

The cells were seeded at a density of 2 x 10^3^ into 96-well plates and treated with increasing concentrations of the drugs for 48 h. Cell viability was determined using WST-1 assay (Sigma). Vehicle-treated cells were used as the control group. For clonogenic survival assay, the cells were seeded onto 6-well plates at a density of 2 × 10^3^ cell/well and incubated at 37 °C in 5% CO_2_ for 14 d. The cultures were then fixed in ice-cold methanol and stained with crystal violet solution (1% w/v). For the clonogenic survival assay, cells were seeded in 6-well plates at a density of 1000 cells/well. After 24 h, the media was changed and the cells were maintained for another 10 d. The resulting colonies were stained/fixed with 0.5% crystal violet imaged using an inverted microscope.

### Drug combination analysis

To explore the efficacy of drug combinations, growth inhibition was determined by WST-1 assay and the results were evaluated by Bliss SynergyFinder ^40^. Visualization of synergy scores is depicted as a synergy map. An average synergy score of 0 is considered additive, less than 0 as antagonistic and over 0 as a synergistic outcome.

### Caspase 3/7 activity assay

The Caspase-Glo® 3/7 Assay System (Promega), a luminescence-based assay for detection of active caspase-3 and 7, was used to quantitatively determine apoptotic cell death. Following caspase cleavage of the proluciferin DEVD substrate, a substrate for luciferase is released and results in luciferase reaction and production of light.

### Western blot analysis

The cells were lysed for 30 min in ice-cold RIPA buffer (50 mM Tris-HCl, pH 8.0, 150 mM NaCl, 1.0% NP-40, 0.5% sodium deoxycholate and 0.1% SDS) containing protease and phosphatase inhibitors (ThermoFisher). The assay was performed with primary and Licor secondary antibodies (Licor Biosciences). β-actin was used as the loading control. The list of antibodies is in Table S5.

### Analysis of gene expression by quantitative reverse transcription-PCR

Quantitative reverse transcription-PCR (qRT-PCR) analysis was done on a QuantStudio Real-Time PCR instrument (ThermoFisher) using PowerUp™ SYBR® Green Master Mix (ThermoFisher). The primer sequences are listed in Table S6. The target gene expression levels were normalized to beta-2-microglobulin (*B2M*) levels. For calculations, 2 ^−ΔΔCT^ formula was used, with ΔΔCT = (CT_Target_ – CT_B2M_) experimental sample – (CT_Target_ – CT_B2M_) control samples, where CT is the cycle threshold.

### Animal studies

The animal experiments were performed according to the Animal Experiment Board in Finland (ELLA) for the care and use of animals under the licenses 4161/04.10.07/2015 and 9241/2018. The animals were kept under pathogen-free conditions in individually ventilated cages in an animal care facility. Mice were kept on a 12 h light/dark cycle with access to autoclaved water and irradiated chow *ad libitum* and were allowed to adapt to the facility for 1 week before starting the experiments. For the subcutaneous experiments, MDA-MB-453 (5 x 10^6^) and HCC1954 (3 x 10^6^) cells were injected into the right flank of six- to eight-week-old BALB/cOlaHsd-Foxn1nu mice (Envigo, France). Mice with tumor size ∼100 mm^3^ were randomized into experimental and control groups. Tumor dimensions were measured with Vernier calipers and tumor volume were calculated as 1/2 larger diameter x (smaller diameter)^2^. For the intracranial model, 1 x 10^5^ cells in 5 µL of PBS were inoculated into the brain of anaesthetized mice. The mice were then imaged with bioluminescence and based on the bioluminescence signal were randomized to experimental and control groups, as descried earlier ^63^. Mice were euthanized when they became moribund when they reached defined study end points.

### Zebrafish studies

Zebrafish embryo xenograft studies were performed under the license ESAVI/9339/04.10.07/2016 (National Animal Experimentation Board, Regional State Administrative Agency for Southern Finland). Briefly, the *DUSP6* knockdown GFP-MDA-MB-361 cells were injected into the brain of zebrafish embryos from the dorsal side. One day after injection (1 dpi), successfully transplanted embryos were placed in CellView glass bottom 96-well plate (1 embryo/well) and embryos were incubated in E3 + PTU at 33 °C. The xenografted embryos were imaged using a Nikon Eclipse Ti2 fluorescence microscope and a 2x Nikon Plan-Apochromat (NA 0.06) objective. Each embryo was imaged at 1dpi and 4dpi using brightfield illumination and a GFP fluorescence filter set (excitation with 470nm LED). Each image was inspected manually to filter out severely malformed, dead or out of focus embryos. The tumor area was measured using ImageJ (NIH). The fold change in tumor size was calculated as follows: GFP intensity (4dpi)/GFP intensity (1dpi)

### RNA-sequencing

RNA-sequencing was conducted at the Finnish Functional Genomic Center, The University of Turku, Finland. RNA was harvested using the NucleoSpin RNA purification kit (Macherey-Nagel), followed by treatment with DNase to remove genomic DNA. RNA (300 ng) was reverse transcribed using the Illumina TruSeq Stranded Total mRNA kit. The quality of the samples was ensured using Agilent Bioanalyzer 2100 or Advanced Analytical Fragment Analyzer. Sample concentration was measured with Qubit® Fluorometric Quantitation (Life Technologies) and/or KAPA Library Quantification kit for Illumina platform, KAPA Biosystems. Sequencing run was performed using the Illumina NovaSeq 6000 instrument. Genomic alignment was performed using Rsubread v. 2.0.0 and the reads were mapped to the human reference genome hg38. Aligned reads were assigned to RefSeq gene models using the same R package with its default settings.

Differentially expressed genes and pathways were identified using the R package limma. Latest hallmark gene sets were downloaded from the Molecular Signatures Database version 7.4 (http://www.gsea-msigdb.org/gsea/msigdb/index.jsp) and used within Gene Set Variation Analysis (GSVA) allowing pathway enrichment estimates for each sample ^64^. Data was transformed using log(x+1) after normalization and the RNA-sequencing pipeline run. Heatmaps were plotted using the ComplexHeatmap package ^65^. R statistical software version 4.0.3 was used for the statistical analyses and visualizations.

### cBioPortal database analyses

The correlation analysis between gene expression and breast cancer subtypes was examined using the METABRIC breast cancer cohort with PAM50 classification^34^, TCGA breast invasive carcinoma dataset and the TCGA Firehose legacy dataset ^35^. The RNA-seq data from breast invasive carcinoma samples and the relevant clinical information are available at the TCGA data portal, cBioPortal for Cancer Genomics (http://www.cbioportal.org/). According to the PAM50 classification, METABRIC breast cancer patients were divided into 5 subtypes, including the basal (n=199), HER2+(n=220), Lum A (n=679), Lum B (n=461) and Normal-like (n=140). ER and HER2 status were assessed using the patient’s IHC information. The survival outcomes were extracted from the TCGA breast invasive carcinoma dataset.

### Statistical analysis

Differentially expressed genes and pathways were identified using the R package limma ^66^. Latest hallmark gene sets were downloaded from the Molecular Signatures Database version 7.4 (http://www.gsea-msigdb.org/gsea/msigdb/index.jsp) and used within Gene Set Variation Analysis (GSVA) allowing pathway enrichment estimates for each sample ^64^, with FDR-cutoff <0.25 used for the pathway _enrichment_ analyses. Data was transformed using log(x+1) after normalization and the RNA-sequencing pipeline run. Heatmaps were plotted using the ComplexHeatmap package ^65^. R statistical software (R Core Team (2023). R: A language and environment for statistical computing. R Foundation for Statistical Computing, Vienna, Austria. URL: https://www.R-project.org) version 4.0.3 was used for the statistical analyses and visualizations.

All data were evaluated in triplicate against the vehicle-treated control cells and collected from three independent experiments. In addition to R analyses, data were visualized and analyzed using GraphPad Prism 8.3.0 using one-way ANOVA and the unpaired two-tailed student’s *t* test. All such data are presented as mean ± standard deviation (SD).

### Reporting summary

Further information on research design is available in the Nature Portfolio Reporting Summary which is linked to this article.

### Data availability

The correlation analysis between gene expression and breast cancer subtypes and the RNA-seq data from breast invasive carcinoma samples plus the relevant clinical information were accessed from cBioPortal ^52^. All other data supporting the findings of this study are available from the corresponding author on reasonable request.

### Code availability

R scripts in this study are available from corresponding author upon request.

## Acknowledgement

Taina Kalevo-Mattila is acknowledged for superior technical support and the entire Turku Bioscience Centre personnel is thanked for excellent working environment. We acknowledge Zhong-Yin Zhang for PRL1i compound, Norma O’Donovan, National Institute for Cellular Biotechnology, Dublin City University, Ireland, for the other HER2+ breast cancer cell lines used in this study, and Dihua Yu, The University of Texas MD Anderson Cancer Center, USA, for the BT474Br cells. We further acknowledge important contributions of the following core facilities of Turku Bioscience Centre (University of Turku and Åbo Akademi University) supported by Biocentre Finland: Finnish Functional Genomics Center, Screening unit, and Zebrafish unit. This study was supported by funding from Finnish Cancer Associations (JW), Foundation of Finnish Cancer Institute (MM), Maud Kuistila Foundation (MM), Turku University Foundation (MM), Finnish Cancer Institute (TDL), Finnish Cultural Foundation (TDL), and Finnish Cultural Foundation (KJK).

## Author contributions

M.M., M.T., D.C., M.S., J.M., A.P., and I.P. performed the experiments. R.V., and T.D.L carried out the data analysis. A.V., K.J.K, and K.E. provided technical and experimental support. M.M., and J.W. designed the study and wrote the manuscript. J.W. supervised the study. All authors read and approved of the manuscript.

**Correspondence and requests for materials** should be addressed to Jukka Westermarck.

---

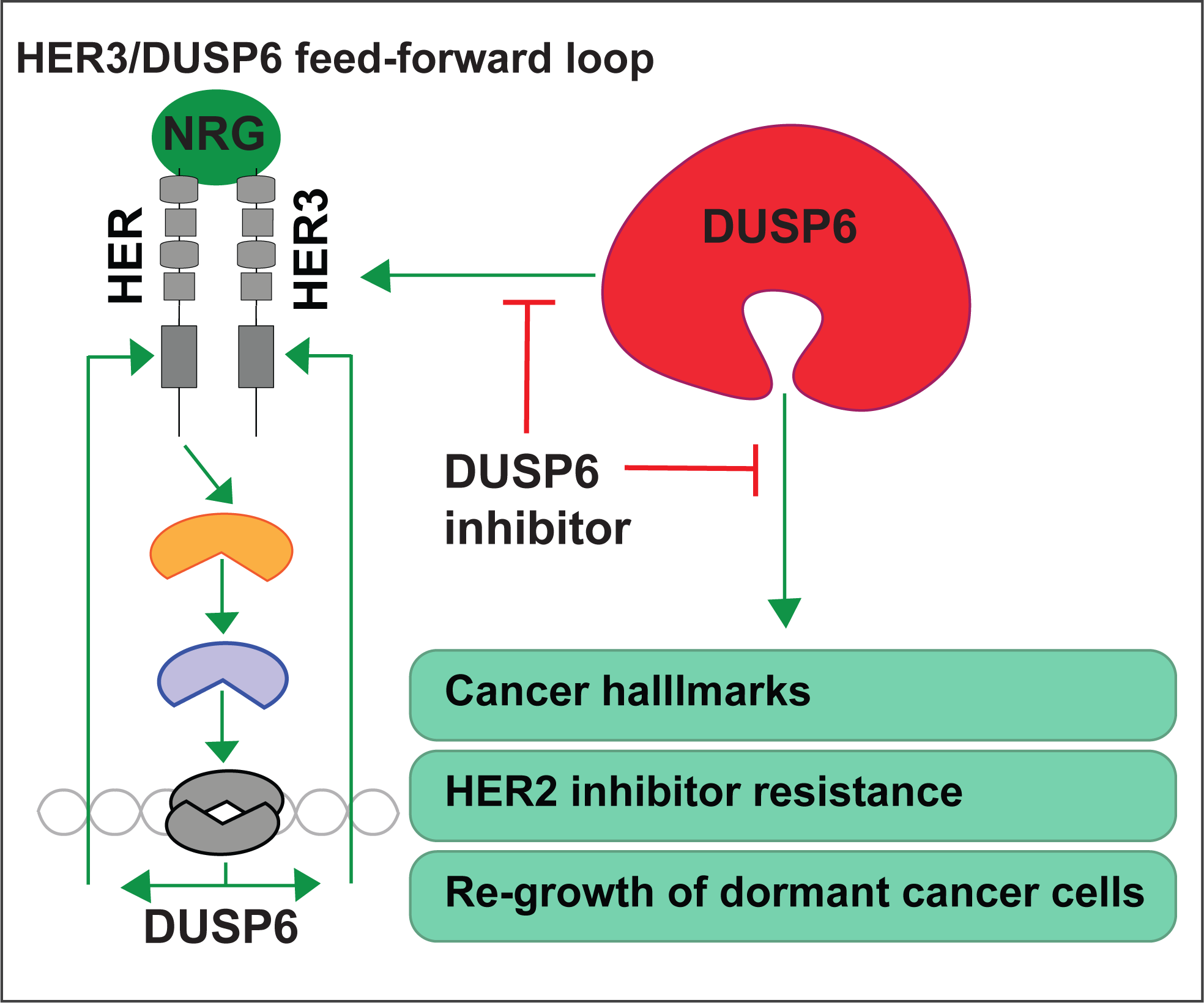

